# Assessing Prediction Performance of Neoadjuvant Chemotherapy Response in Bladder Cancer

**DOI:** 10.1101/075705

**Authors:** Chris Cremer

## Abstract

Neoadjuvant chemotherapy is a treatment routinely prescribed to patients diagnosed with muscle-invasive bladder cancer. Unfortunately, not all patients are responsive to this treatment and would greatly benefit from an accurate prediction of their expected response to chemotherapy. In this project, I attempt to develop a model that will predict response using tumour microarray data. I show that using my dataset, every method is insufficient at accurately classifying responders and non-responders.

## I. INTRODUCTION

Bladder tumours are classified based on their stage and grade [1]. Grade is given based on the cancer cells’ appearance compared to normal cells and pathological stage (pT) describes the extent of penetration of the tumour. These two clinical features are currently important factors when deciding which treatment to administer to patients with bladder cancer.

Neoadjuvant chemotherapy is the administration of chemotherapeutic agents before a standard treatment. The current standard treatment for muscle-invasive bladder cancer (MIBC) is radical cystectomy, which is the surgical removal of all or part of the bladder. The rationale for the administration of chemotherapy prior to cystectomy is that it can help treat micrometastatic diseases and downstage the diagnosed tumour so that there is an increase potential for complete resection of the tumour [2]. It has been shown that compared to radical cystectomy alone, the use of neoadjuvant chemotherapy followed by radical cystectomy increases the likelihood of eliminating residual cancer in the cystectomy specimen and is associated with improved survival among patients with locally advanced bladder cancer [3]. However, the overall response rate of neoadjuvant chemotherapy is only 50% to 60% [4]. Consequently, neoadjuvant chemotherapy negatively affects the health of non-responsive patients while they receive no benefit from the harmful treatment. Therefore, there is a critical need to predict those who will respond positively to neoadjuvant chemotherapy.

The expression of some genes have been found to have a correlation to chemotherapy response. BRCA1 plays a central role in DNA repair pathways. Font et al. found that patients with low/intermediate BRCA1 levels had a higher pathological response rate than those with high levels [5]. Similarly, Kiss et al. observed that the overexpression of Bcl-2, an inhibitor of the apoptotic cascade, in chemotherapy-naive primary tumors is related to poor response to neoadjuvant chemotherapy, which might help discriminate likely non-responders [6]. The expression of genes, such as BRCA1 and Bcl-2, are examples of features that can be used to distinguish responder vs non-responder.

Furthermore, muscle-invasive bladder cancers are biologically heterogeneous and can be stratified into 2-4 subtypes, such as basal, luminal, and p53-like [7]. These subtypes can have widely variable clinical outcomes and responses to conventional chemotherapy [8]. Specifically, p53-like MIBC are consistently resistant to neoadjuvant MVAC chemotherapy, and all chemoresistant tumors adopt a p53-like phenotype after therapy. Therefore, identifying subtypes can be another useful approach to predicting response to chemotherapy.

Various groups have come up with different methods to predict patient response to chemotherapy. One approach uses a method called coexpression extrapolation (COXEN) derived from expression microarray data of the National Cancer Institute (NCI)-60 cell line panel to predict drug sensitivity of bladder cancer cell lines [9], [10]. To test its performance, they performed in vitro drug response experiments to determine the sensitivity of each bladder cell line to cisplatin and paclitaxel. For those cells, prediction accuracies averaged 85% for cisplatin and 78% for paclitaxel. A different group study by Takata et al. predicted response to chemotherapy by selecting genes that discriminated responder vs non-responder then scored each sample by a weighted-vote of each gene [11]. In a validation study, they applied their method to 22 additional cases of bladder cancer patients and found that the scoring system correctly predicted clinical response for 19 of the 22 test cases [12]. In 2011, the same group applied their method to the same problem that this project faces: predicting response of gemcitabine and cisplatin treated muscle-invasive bladder cancer using microarray data. They achieved strong results, correctly classifying 18 out of 19 test samples [13].

There are numerous machine learning classification methods that can be applied to this problem. For instance, logistic regression is a method that defines a linear decision boundary by minimizing an error function of the training samples’ target and output. In contrast, k-Nearest Neighbors is a method that decides a sample’s class based on the class of the k samples that are nearest to it. Consequently, depending on the method, the predicted class can be determined very differently.

In this project, I apply several different classification methods to bladder tumour microarray data and try to predict response. I also compare the performance of these methods to the method developed by Takata et al. by applying them to the same dataset. Finally, I attempt to use a neural network to model the complex features in the data with hopes of improving prediction accuracy.

## II. MATERIALS AND METHODS

### A. Microarray Data

The data used to predict response to chemotherapy is microarray data (17000 gene expressions) for 52 muscle-invasive bladder cancer patients who received gemcitabine and cisplatin chemotherapy after their initial biopsy. Out of the 52 patients, 15 (29%) were complete responders (pT<2 and lymph node negative) and 37 (71%) were non-responders (pT2 or higher, or any node positive). The data was log-transformed and quantile normalized with R package aroma.light. Prior to analyzing the data, a minimum cutoff mean and standard deviation gene expression was set to remove genes that had either very low expression or had expressions that did not significantly vary. This helped reduce the dimensionality of the data to nearly 9000 genes.

### B. Nested Two-Fold Cross-Validation

Since there is only 52 samples to use for training, validation, and testing, I decided to implement nested two-fold cross-validation. Nested two-fold cross-validation means that the data is split in two, one half for training and the other half for testing, and then the training half is split again. The first fold is used for testing the performance of the classifier on held out data and the inner fold is used to select hyperparameters for the models. More specifically, I use stratified two-fold cross-validation so that the proportion of each class remains relatively constant for each fold.

### C. Machine Learning Methods

The following are the classification methods used in this project which were imported from Scikit Learn: Logistic Regression with L1 (LR1), L2 (LR2), and ElasticNet (LRel) regularization, k-Nearest Neighbors (kNN), Random Forests (RF), Support Vector Classification (SVC), Gaussian Naive Bayes (GNB), Linear Discriminate Analysis (LDA) and AdaBoost (ADA). These methods were chosen because they exhibit a wide range of different decision boundaries given the same data set. There hyperparameters were set by two-fold cross-validation.

### D. Takata/Kato 2011 Method

I implemented the method used by Kato et al. [13] which was shown to be an effective method at predicting response given the same scenario. Their method first identifies genes that discriminate between the two classes then calculates a prediction score (PS) for each test sample using those discriminating genes. Mean (µ) and standard deviation (δ) were calculated from the log-transformed data of each gene in the responder (R) and non-responder (N) cases. A discrimination score (DS) for each gene was defined as:

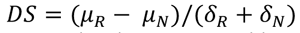

Next, samples were randomly permutated between the two classes 10,000 times and since the DS data set of each gene showed a normal distribution, the P-value for each gene was calculated. Cross-validation is used to determine how many of the top significant genes will be used in the gene set. To classify a sample, each gene in the gene set votes for either responder or non-responder depending on whether the expression level (x_i_) in the sample is closer to the mean expression level of the responders or non-responders in the reference samples. The magnitude of the vote (V_i_) reflects the deviation of the expression level in the sample from the average of the two classes.

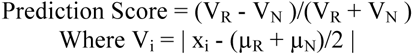

Therefore, a positive PS predicts responder whereas a negative PS predicts non-responder and a higher absolute value reflects a stronger confidence in the prediction.

### E. Neural Network

To implement a neural network, I modified the Python code from http://neuralnetworksanddeeplearning.com/ to fit my problem. The network had sigmoid activation functions for each layer and cross-entropy error. The hyperparameters, such as network topology, learning rate, and regularization were determined through cross-validation.

### F. Accuracy vs AUC

Since there is an imbalance in the number of each class (15 responders vs 37 non-responders), if we predict all samples to be non-responders we will achieve 71% accuracy and only 50% AUC. I have chosen to report the AUC in Fig 1 and 2 instead of accuracy for each method because it is unaffected by the imbalanced classes and in this case is a better indicator of a method’s predictive performance.

**Fig 1:**
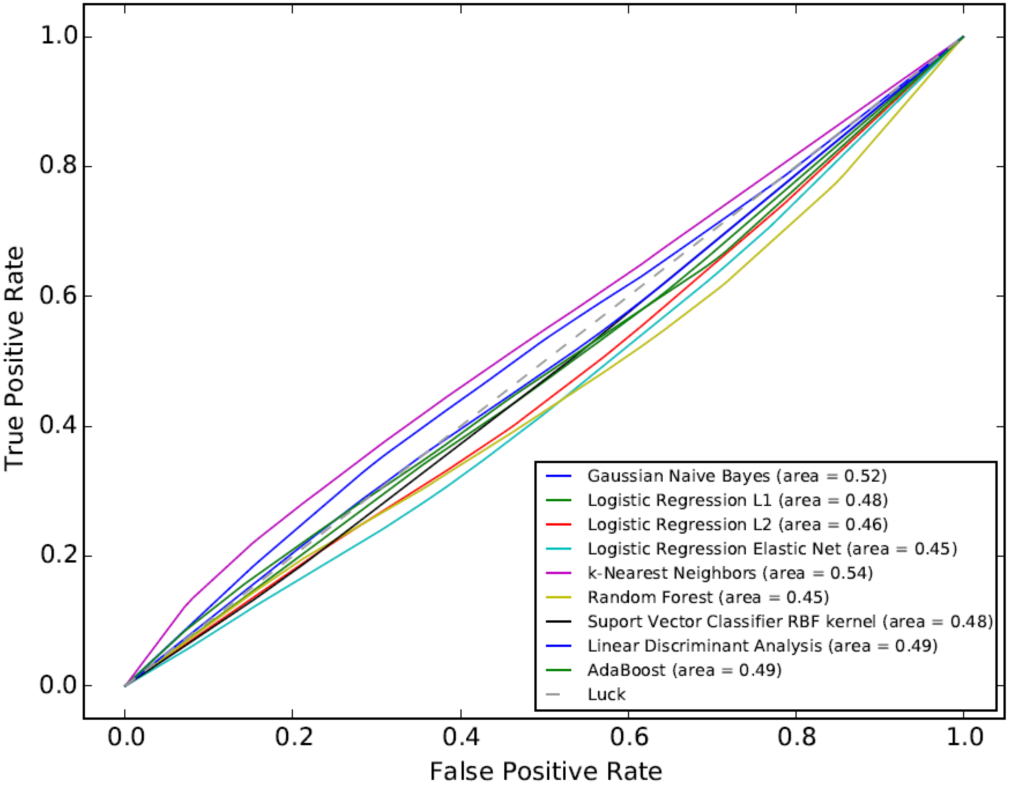
True Positive Rate vs False Positive Rate of several classification methods based on averaging 10 iterations. All methods have an AUC around .5

**Fig 2:**
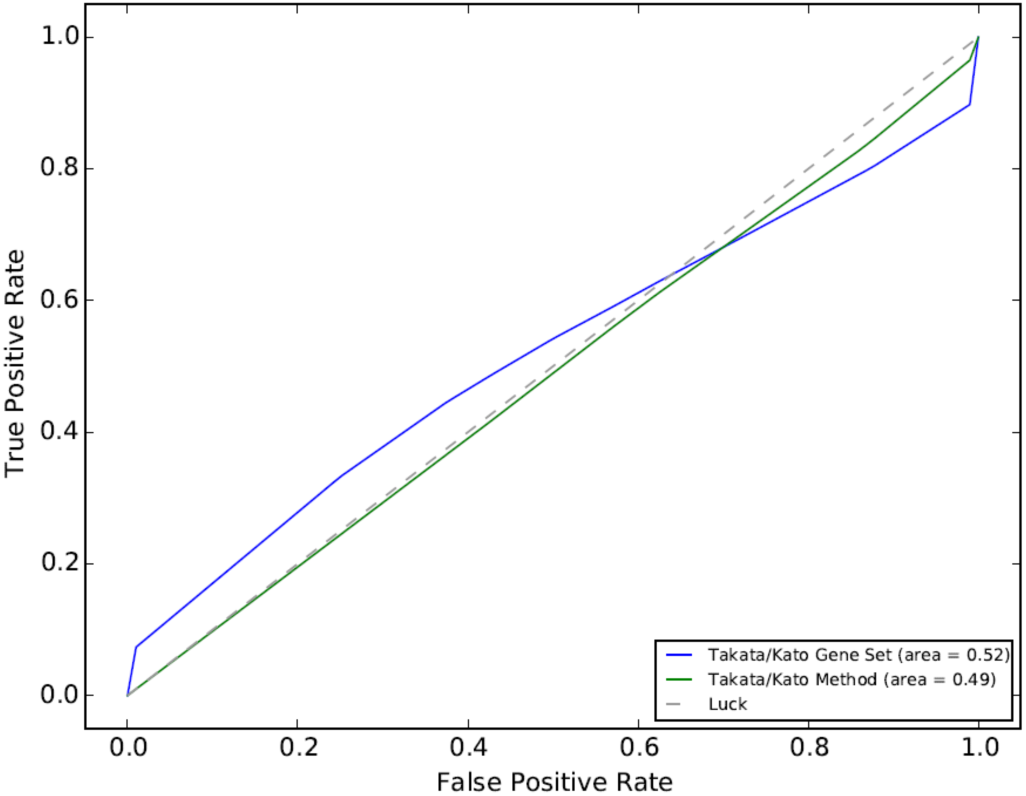
True Positive Rate vs False Positive Rate of the method from [13] using the gene set they published (blue line) as well as the gene set obtained from applying their published method to my data. Both gene sets have an AUC of about 0.5 (as good as random).

## III. RESULTS

### A. Prediction Results

I applied various classification methods to the microarray data. Fig. 1 is a plot of the average receiver operating characteristic (ROC) curves over 10 iterations of these various methods. The methods outputted 0 for responders and 1 for non-responders. The plot shows that none of the methods are capable of properly classifying the two classes. The AUC of methods are all near 50%, indicating they are essentially random. The highest AUC is kNN at 0.54 and the lowest is Random Forest at 0.45. Table 1 shows the accuracy of the methods. Most methods learn to achieve 71% by predicting non-responder for all samples which is about .50 AUC. The continuous value output of most methods is about 0.71 (ie. predicts non-responder), which is equal to the average label value (37/52). This output is due to the imbalanced classes (15 responders vs 37 non-responders) because it minimizes the error when classes are predicted by chance. If the classes are balanced by subsampling the majority class, then the methods output 0.5 for most samples, indicating they are unable to predict class.

**Table 1.** Training and validation accuracies and AUC of various methods predicting responder vs non-responder using microarray data. The validation accuracies are computed from leave-one-out cross-validation. There are 52 samples (15 responders, 37 non-responders).

| Method | Training Accuracy (%) | Validation Accuracy (%) | Validation AUC (%) |
| --- | --- | --- | --- |
| Gaussian Naive Bayes | 100 | 71 | 52 |
| Linear Discriminate Analysis | 77 | 71 | 49 |
| Support Vector Classification | 71 | 71 | 48 |
| k-Nearest Neighbors | 73 | 67 | 54 |
| Logistic Regression with L1 Reg. | 100 | 63 | 48 |
| Logistic Regression with L2 Reg. | 100 | 63 | 46 |
| Logistic Regression with ElasticNet | 71 | 71 | 45 |
| Random Forests | 98 | 63 | 45 |
| Takata Method | 92 | 69 | 49 |
| Takata Gene Set | 64 | 48 | 52 |

### B. Kato Method Results

Since the various classification methods I used previously were not successful at distinguishing the two classes, I implemented the method from [13] which was shown to be an effective method given the same classification problem. See ‘Kato 2011 Method’ in Material and Methods for a description of the method. In their paper they came up with a gene set that could correctly classify 18 out of 19 test samples. I applied their method to my data to come up with another gene set. Using the gene set obtained from their data and the one from my data, I calculated a prediction score for each test sample in my data set. Figure 2 is the ROC curve of the predictions using each gene set. The AUC is nearly .50 for both, which is in indication that their predictions are essentially random. This is the same result as Fig 1; the genes in the training set that are found to be significant to not generalize to the test set.

### C. Neural Network Results

In an effort to identify more complex features in the data to improve predictions, I implemented a neural network. I experimented with a wide range of hyperparameters. For instance, I tested networks with 1 to 3 hidden layers and 2 to 1000 nodes per layer. Fig. 3 is a comparison of the error on the training set vs the test set over 300 epochs of a network with a topology of two hidden layers with 10 nodes each. Most network topologies resulted in a very similar plot as Fig 3. The training error is seen to descend initially then level off until epoch 200 where it begins to descend again. In contrast, the test set error increases initially then fluctuates at a high error for the remaining epochs. This is an indication that the features learned in the training set do not generalize to the test set. If the problem with the model was overfitting, we would expect the test set error to initially drop then begin to increase when the model overfits. If the model were underfitting, then we would expect the training error to not descend at higher epochs. Thus it is not a problem of overfitting or underfitting, it’s that there is not enough structure in the data to make strong predictions. Fig 5 in the Supplemental section is a comparison of the network trained on the real microarray data and the network trained on data that is completely random. The random data was created by sampling from a uniform distribution for each feature of each sample. The two graphs are similar in that the test error does not decrease when the training error decreases, but they are different at the later epochs as the real data graph A) has high fluctuations in its test error whereas the simulated random data does not. Nonetheless, Fig 5 is an indication that the real data may not have much more structure than random data.

**Fig 3:**
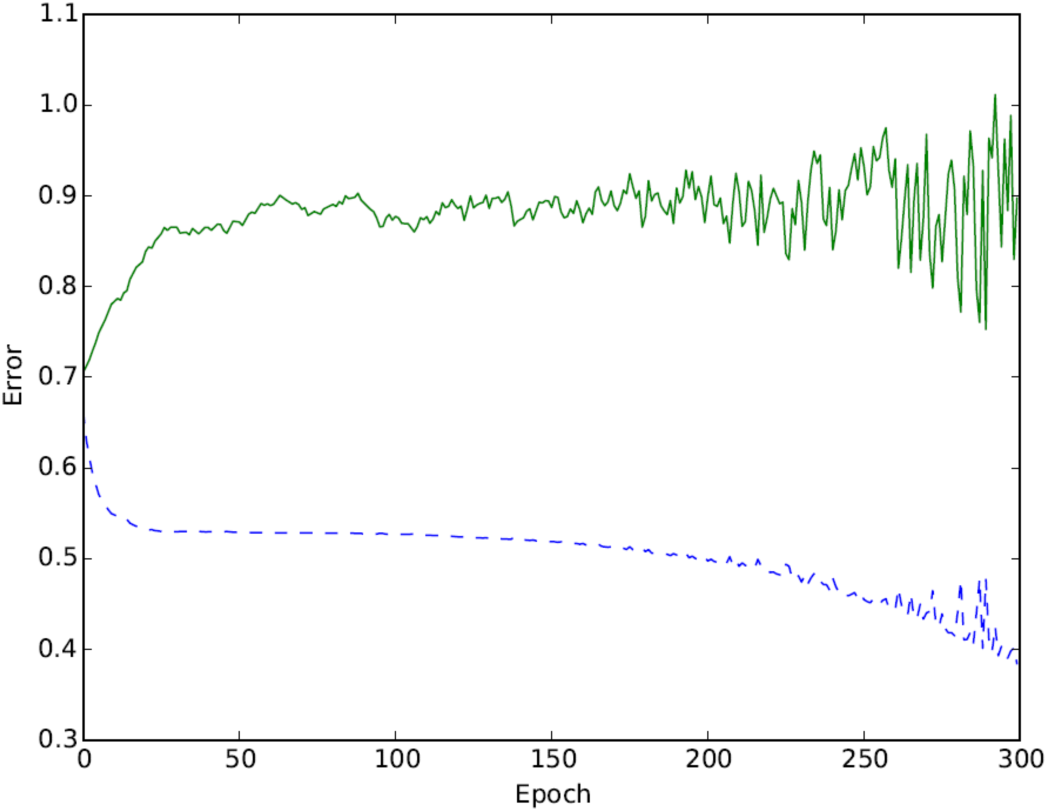
Neural Network Training (dashed blue) and Test Error (solid green) over 300 epochs.

**Fig 5:**
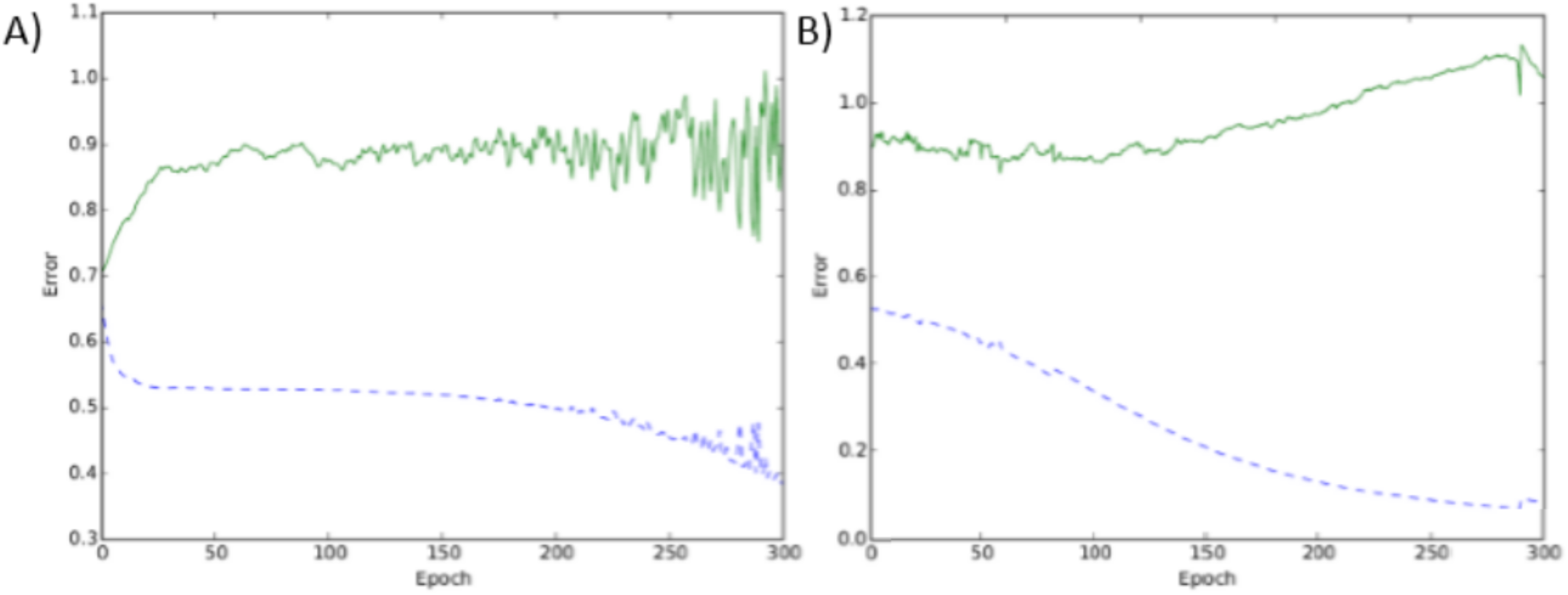
A) Neural Network Training (dashed blue) and Test Error (solid green) over 300 epochs on microarray data. B) Same network applied to random data for comparison.

## IV. DISCUSSION

### A. Possible Explanations for Poor Results

Takata and Kato have shown previously that their method could discriminate responders and non-responders almost perfectly (18/19 correct test samples). However, using my dataset, they achieve an AUC of nearly 0.5, meaning their method was effectively random, just like the results of all the other methods I tried. As an illustration of the difference between datasets, Fig 4A is a figure from Kato’s paper [13] showing the prediction scores of the training and test sets. It is clear that the scores are clearly distinct between the two classes. Fig 4B is the prediction scores of the test set from applying the same method to my dataset. There is no longer any difference between the responders and non-responders. The reason for this outcome could be that the measurements of the gene expressions from the microarrays in my dataset may have been very noisy causing the input data to be imprecise. There could have been some sort of batch effect or preprocessing that may have modified the negatively modified the inputs. Similarly, there could have been errors when labeling the samples as responders or non-responders. For these possible reasons, the method used by Takata and Kato performed much better on their dataset than mine.

**Fig 4:**
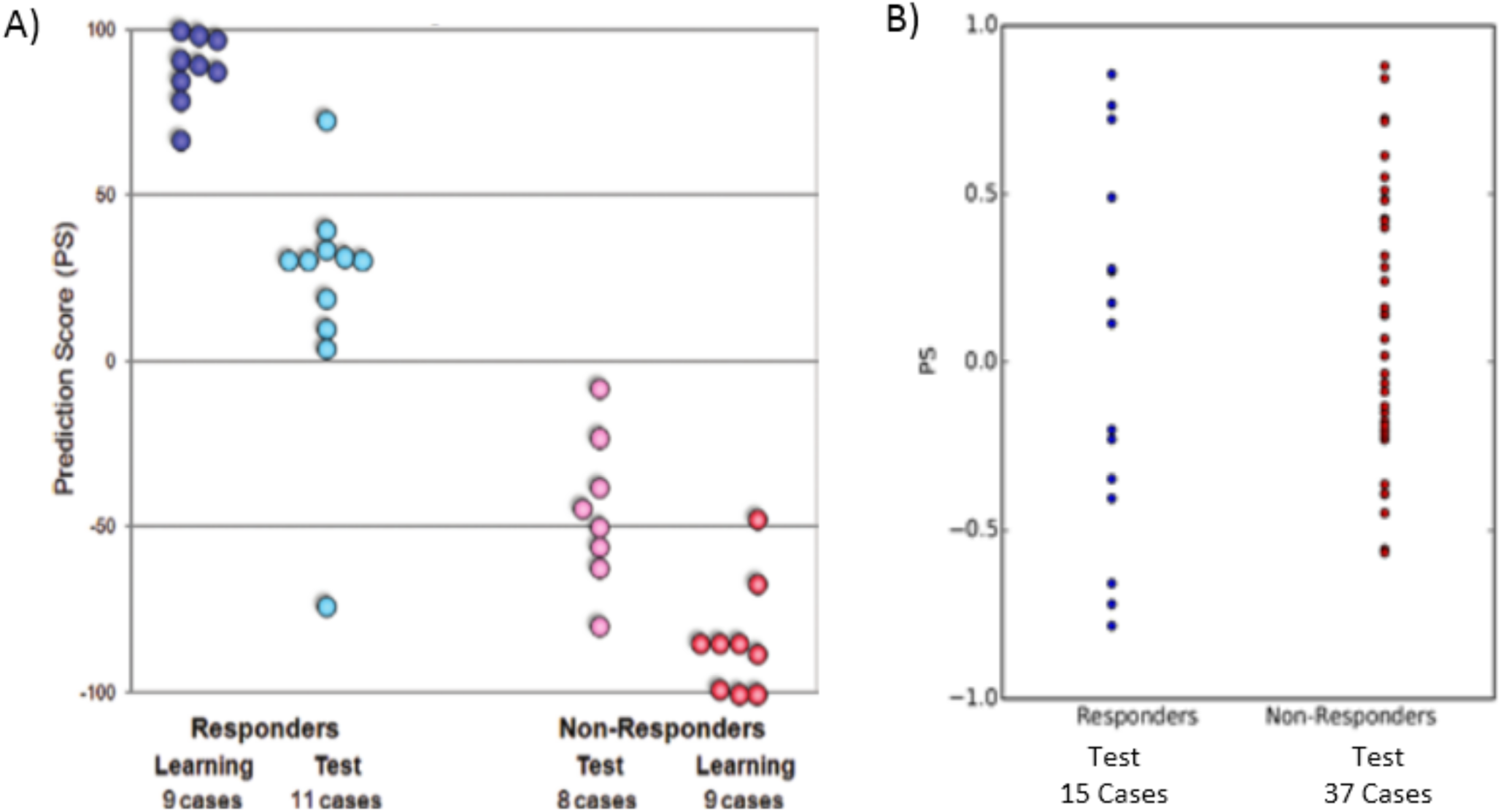
*A) Figure from* [13] *showing the distribution of the prediction scores between classes for training samples and test samples. Test samples were predicted correct 18/19. B) The prediction scores of the test samples using the same method applied to my dataset. There is effectively no distinction between classes. I subsampled from the majority class (non-responders) so that the predictions would not be biased.*

### B. Relabeling Data

To see if the reason for the poor accuracy was due to mislabeling, I tried relabeling the data based on different definitions of being a responder. Initially, the definition of responder was that the post-chemo sample be at a stage less than 2 (pT<2) and be lymph node negative. One example of a new responder definition is if there is any decrease in stage of the tumour (not necessarily less than 2) then we classify that sample as a responder. I tested this relabeling scheme and others but unfortunately the methods were still unsuccessful at effectively classifying the two classes.

### C. Simulated Data and L1 Logistic Regression

As a proof of concept, I simulated data to compare the effectiveness of the various methods. The assumption I make for the simulated data is that the majority of the features/genes are non-informative for distinguishing the two classes but there is a small subset of the features that are informative. So for the majority of the features, I sample from the same Gaussian for both classes, whereas for a minority of the features, each class is sampled from two different (possibly overlapping) Gaussians. If I make 100 samples (50 for training, 50 for validation) with 100 features and 10 of those features being distinguishable, then most of the methods perform well (>96% accuracy). If I increase the number of features to 9000 (similar to this project) and keep 10 distinguishable features, then most methods do poorly (<60% accuracy) except Logistic Regression with L1 regularization (>90% accuracy). This must be because L1 gives zero weight to all features that don't seem useful whereas the other methods give a little weight to most features which ends up being a significant amount of noise when predicting validation samples. This result is in accordance with Andrew Ng’s paper on L1 regularization and irrelevant features [14]. Basically, the simulated data showed that if there were genes that were differentially expressed then atleast L1 Logistic regression should be able to find them.

### D. Other Attempts

I tried taking the majority vote of all methods for each sample to decide its class. The result was that the combination of methods was either the same or worse than any individual method.

Next, rather than trying to find significant genes, I investigated whether genes that have been reported to have an influence on chemotherapy response were predictive in my dataset. To this end, I selected genes BRCA1, Bcl-2, and p53 and ran my methods on them individually. None of the methods achieved an accuracy better than using all genes in the dataset, meaning that these specific genes were not predictive in this case. Finally, instead of applying supervised methods, I tried clustering the data to see if the assigned labels would have any relation to response. I first applied PCA to reduce the dimensionality to 50 then fitted a Gaussian Mixture Model with two clusters. The predicted classes ended up not having any sensible relation to the response of the samples.

### E. Future Directions

For my future directions, I will aim to acquire a new dataset. Ideally, the new dataset will have more samples which will reduce the overfitting of my models and reduce the effect of outliers. I can also try using RNASeq data instead of microarray data. The TCGA would be a good source for this data. Furthermore, rather than predicting from single gene expressions, we could look at pathways. We could create metagenes based on the expressions of genes involved in the same pathway. These metagenes would help reduce dimensionality, increase regularization, and help the interpretability of the model. However, identifying these metagenes would not be trivial and would be an area of future work. Additionally, I will be incorporating mutational data into my analyses to potentially help the predictions.

## V. CONCLUSION

The goal of this project was to improve the prediction of response of muscle-invasive bladder cancer to neoadjuvant chemotherapy. Given my dataset, none of the methods that I used were able to effectively distinguish responders from non-responders. In addition, the method used in Kato et al. was also unable to accurately classify the two classes using my dataset. Furthermore, the features learned by the neural net using the training set did not generalize to the test set. My negative results may stem from the dataset being too noisy to accurately predict response. Future work should incorporate data from other sources, such as the TCGA.

